# Synergistic effects of inhibitors targeting PI3K and Aurora Kinase A in preclinical inflammatory breast cancer models

**DOI:** 10.1101/2023.08.11.552992

**Authors:** Nadia Al Ali, Jacob Kment, Stephanie Young, Andrew W.B. Craig

**Affiliations:** Department of Biomedical and Molecular Sciences, Queen’s University, Kingston, Ontario, Canada; Division of Cancer Biology & Genetics, Queen’s Cancer Research Institute, Kingston, Ontario, Canada

**Keywords:** Inflammatory Breast Cancer, Targeted therapy, Combination therapy, PI3K inhibitor, Aurora Kinase A inhibitor

## Abstract

**Background:** Inflammatory breast cancer (IBC) is an aggressive clinical subtype of breast cancer often diagnosed in young women. Lymph node and distant metastases are frequently detected at diagnosis of IBC, and improvements in systemic therapies are needed. For IBC that lack hormone or HER2 expression, no targeted therapies are available. Since the phosphatidyl inositol 3’ kinase (PI3K) pathway is frequently deregulated in IBC, some studies have tested the pan PI3K inhibitor Buparlisib (BKM120). Although the SUM149 IBC cell line was resistant to Buparlisib, a functional genomic screen showed that silencing of Aurora kinase A (AURKA) sensitized cells to killing by Buparlisib. In this study, we tested whether combination treatments of PI3K and AURKA inhibitors act synergistically to kill IBC cells and tumors.

**Methods:** SUM149 cells were treated with increasing doses of PI3K inhibitor Buparlisib (BKM120) and AURKA inhibitor Alisertib as monotherapies or combination therapies. Effects on target pathways, cytotoxicity, cell cycle, soft agar colony growth and cell migration were analyzed. The individual and combined treatments were also tested in a mammary orthotopic SUM149 tumor xenograft model to measure effects on tumor growth and metastasis

**Results:** The SUM149 IBC cell line treated with Buparlisib showed reduced PI3K/AKT activation but no significant skewing of cell cycle progression. Parallel studies of Alisertib treatment showed that AURKA inhibition led to a significant block in G2/M transition in SUM149 cells. In cytotoxicity assays, Buparlisib and Alisertib combination treatments were highly synergistic compared to monotherapy controls. Evidence of synergy between Buparlisib and Alisertib also extended to soft agar colony growth and wound healing motility in SUM149 cells. The combination of Buparlisib and Alisertib also reduced IBC tumor growth in mammary orthotopic xenograft assays and reduced spontaneous metastases development in lung tissue.

**Conclusions:** Although SUM149 IBC cells were relatively resistant to killing by the PI3K inhibitor Buparlisib, our study showed that co-targeting the mitotic kinase AURKA with Alisertib synergized to limit IBC cell growth and motility, as well as IBC tumor growth and metastasis.

## Background

Inflammatory breast cancer (IBC) is an aggressive clinical subtype of breast cancer that often presents as breast swelling and redness of the breast’s skin (1, 2). IBC is frequently diagnosed in young women and often at later stages, with metastases to lymph nodes or distant sites (1). IBC represents ~6% of all breast cancer cases worldwide (3). When categorized by molecular subtypes, 12% of IBC tumors are triple-negative (lacking ER/PR/HER2) breast cancers (TNBC). TNBC have high metastatic capabilities and worse prognosis compared to other subtypes with more options for targeted therapies (1).

Driver mutations of IBC remain unclear, however a study profiling somatic mutations in IBC tumors revealed high heterogeneity and high mutational burden compared to non-IBC breast tumors (4). The three most frequent pathways altered in IBC were PI3K/AKT, Ras/MAPK, and cell cycle pathways (4). The Phosphoinositide 3’ kinase (PI3K) pathway regulates cell growth, survival, metabolism, motility, and angiogenesis (5). Considerable efforts have been underway to target aberrant PI3K activity in breast cancer with direct PI3K inhibitors, or inhibitors of effectors AKT and mTOR kinases (5, 6). Limited responses to monotherapies and acquired drug resistance must be overcome to optimize treatments targeting the PI3K pathway (5, 7).

A recent study investigating resistance mechanisms in IBC and non-IBC models treated with pan PI3K inhibitor Buparlisib reported that gene silencing of several protein kinases, from a kinome-wide screen, overcame resistance in breast cancer cell lines (8). The authors validated MEK1 & PI3K synthetic lethality across breast cancer models with both gene silencing and Selumetinib/Buparlisib combination treatments in cell lines and brain metastatic mouse models (8). Another gene silencing hit that sensitized SUM149 IBC cells to killing by Buparlisib was Aurora Kinase A (AURKA), but this synthetic lethal interaction was not validated (8). AURKA is a nuclear serine/threonine kinase that is activated in G2 phase of the cell cycle and regulates cell division (9). Specifically, AURKA regulates centrosome maturation, entry to mitosis and assembly of the mitotic spindle (10). AURKA overexpression and gene amplification occurs in multiple cancers with links to poor prognosis and increased genomic instability (11). Alisertib (ALS) is an orally available AURKA inhibitor (12), and is one of several AURKA inhibitors to be tested in early phase clinical trials for several cancer types, including hormone receptor-positive breast cancer (9, 13).

In this study, we investigate whether combination treatments of PI3K and AURKA inhibitors can limit IBC cell growth, viability and motility in 2D and 3D cell culture models. Buparlisib and Alisertib were also tested as both monotherapies and in combination in mammary orthotopic SUM149 tumor xenograft models. In most of the above assays, the combination of Buparlisib and Alisertib showed synergistic effects in limiting the IBC cell/colony growth and motility, and tumor growth *in vivo*.

## Methods

### IBC cell line and media

The human SUM149PT cell line was isolated from an IBC tumor and purchased from a commercial source (BIOIVT). SUM149 cells were cultured in Ham’s F-12 media supplemented with antibiotic-antimycotic (1%), and 5% fetal bovine serum (FBS). SUM149 cells were routinely tested for mycoplasma contamination and were confirmed mycoplasma free.

### Drug preparations

Buparlisib and Alisertib were purchased from Med Chem Express (MCE), and solubilized in dimethyl sulfoxide (DMSO) for *in vitro* assays. For *in vivo* studies, Buparlisib and Alisertibe were solubilized stepwise in 10% DMSO, 40% PEG300, 5% Tween-80 and 45% sterile saline.

### Cell synchronization, lysates and immunoblotting

For testing effects of inhibitors on PI3K/AKT pathway, lysates were prepared from subconfluent SUM149 cells treated with a vehicle control (DMSO), Buparlisib (5 μM), Alisertib (5 μM) or a combination of both drugs at the same concentration for 1 hour. To measure the effects of the inhibitors on pAURKA/AURKA protein levels, we performed a double thymidine block and release to synchronize SUM149 cells prior to preparing cell lysates. Briefly, subconfluent SUM149 cells were treated with thymidine at a final concentration of 2 mM for 19 hours. Following a rinse with PBS, cells were allowed to recover for 9 hours in fresh medium prior to a second round of thymidine (2 mM) treatment for 16 hours at 37 °C. Cells were released from G1/S block by washing with prewarmed PBS and incubated in fresh media for 5 hours before adding DMSO, Buparlisib (5 μM), Alisertib (5 μM) or a combination of both drugs. Cells were collected at 0, 5, 8, 10, and 12 hours of treatment for flow cytometry analysis of cell cycle by DNA content following DAPI staining (CytoFLEX, Beckmann). Cell lysates were analyzed by immunoblotting with pan AURKA and phospho-AURKA (pAURKA) antibodies (CST). For both methods, cells were lysed on ice with supplemented RIPA buffer (50 mM Tris, 5 mM EDTA, 150 mM NaCl, 1% NP-40, 0.5% Sodium Deoxycholate, 0.1% SDS, 10 μg/mL aprotinin, 10 μg/mL leupeptin, 1 mM Na_3_VO_4_,100 μM phenylmethylsulfonyl fluoride, 50 mM NaF) and centrifuged at 3000 × g for 15 min at 4°C. Supernatants were collected and proteins concentrations were measured with Bradford assay. An amount of 100 µl sample was resolved by sodium dodecyl sulfate polyacrylamide gel electrophoresis (SDS-PAGE) sample loading buffer and electrophoresed on 10% SDS-PAGE gel after thermal denaturation at 95°C for 5 min. Samples were transferred onto PVDF membrane at 25 mA for 15 min at RT (TurboBlot, BioRad). After blocking, the membranes were probed with indicated primary antibodies from CST overnight at 4°C: pan AKT (1:1000), phosphoS274-AKT (1:1000), pan AURKA (1:1000), and phosphoT288-AURKA (1:1000), all primary antibodies were anti-rabbit except for the loading control B-actin, anti-mouse. After washing, fluorophore-conjugated secondary antibodies (Goat anti-Rabbit IgG (H+L) Dylight™ 800 4X PEG and Goat anti Mouse IgG (H+L) Dylight™ 680 secondary antibodies from Invitrogen, Thermofisher) were incubated for 1 hour at room temperature prior to detection and quantification using an Odyssey CLx scanner (LI-COR).

### Cell cycle analysis

SUM149 cells (10^6^) were incubated with drugs for 48 hours at Buparlisib or Alisertib 5 μM, and same for combination. At the end of the 48 hours, cells were treated according to the Abcam protocol. Cells were harvested and washed in PBS, then fixed in 70% cold ethanol by adding drop wise to pellet while on vortex. Cooled at 4°C and centrifuged and the supernatants were discarded carefully. Treated with ribonuclease and PI respectively then cell cycle distribution was analyzed by flow cytometry (FACS Aria III, BD). Analysis was done using the Flow Jo flow cytometry analysis software.

### Cell viability assay

SUM-149 cells were seeded at 1 x 10⁴ cells per well in a 96-well plate overnight prior to addition of vehicle (DMSO), Buparlisib or Alisertib alone (0.1, 1 or 10 μM), or in combinations at the different doses. Alamar Blue 10% was added to each well and incubated for 4 hours under standard conditions of 37 °C and 5% CO2 in a humidified atmosphere and absorbance at 570 nm (600 nm reference wavelength) using a plate reading spectrometer at 48 hours. All analyses were performed using graphing and statistic packages in GraphPad Prism was used for calculations and statistics. SynergyFinder Bliss model was also used to calculate and present the interaction between Buparlisib and Alisertib in vitro with a dose matrix test. Graphs and statistics were produced by the SynergyFinder application (14).

### Soft agar assay

SUM149 cells were plated at low density in 6-well plates and cultured for 21 days in low melt agar with media + drugs (Buparlisib 2.5 μM, Alisertib 5 μM and combination of 2.5 + 5 μM) changed every 2 days depending on half-life of each drug. At the end of the culture days, the colonies were stained with 5 % crystal violet for 1 hour then washed. The images of the colonies were captured using QCapture pro software attached to a camera on a dissection microscope. Each assay was performed in triplicate, plates were maintained at 37C under 5% CO2 for three weeks. Colonies greater or equal to 100 μm in diameter were counted, as described previously (15).

### Wound healing migration assay

SUM149 cells were seeded at a density of 3.0 × 10^5^ cells per well in 96-well culture plates overnight so that the cells would attach. A single wound was made on each well for each cell line by scratching the attached cells using ESSEN Bioscience Wound Maker 96. The plates were washed with PBS to remove cellular debris from the scraped surface. Drugs were added to SUM149 cells (Buparlisib 1 μM, Alisertib 1 μM and combination 1+1 μM). Cells were incubated in Incucyte Zoom (Sartorius) for 24 hours and the images of the cells were taken immediately and every 2 hours using Incucyte Zoom with a 10X objective.

### Mammary orthotopic IBC tumor xenograft model

Mammary orthotopic tumor xenograft assays were performed using SUM149 cell model and Rag2^−/−^:IL2Rγc^−/−^ recipient female mice, lacking natural killer, B, and T cells. Animals were housed in a specific pathogen-free facility (Queen’s University Animal Care Services), with ventilated cages and sterilized food and water supply. All procedures with mice were approved by the Queen’s University Animal Care Committee. Cells were grown to 70–85% confluence before trypsinization and counting. For xenografting, 5 × 10^5^ SUM149 cells (transduced with WPI-Luciferase) were injected into the right thoracic mammary fat pads of the Rag2^−/−^:IL2Rγc^−/−^ female mice in a volume of 100 μl containing 50% Matrigel using a hypodermic syringe. Tumours were screened to have average initiating size. Mice were grouped into 4 groups each with 6 individuals. One group received vehicle, the other two, one received Buparlisib and the other Alisertib, and the last received the combination. Drugs and doses were based on previous conducted studies (25 mg/kg) per each drug and per combination. Treatment started on day 10 post injection for a (5+2) regimen, at end points 5 weeks, mice were killed, and primary tumor mass recorded. Several tissues were removed for detection of metastases, which were primarily observed in the lungs. The primary tumors and lungs from each mouse were used for histological analysis. For lung inflation, use a 3mL syringe with a 22g needle, this time held parallel to the trachea, insert the needle into the trachea and inject 10% formalin with rate of flow no greater than ∼200 µL/second until the lungs have fully inflated. Once the lungs are inflated, formalin will backflow out of the trachea. Samples were fixed in formalin and embedded in paraffin, and 5 μm sections were stained with hematoxylin and eosin.

### Statistical Analysis

Statistical analyses were performed in GraphPad Prism (version 9.4.1, GraphPad Software). * = p < 0.05, ** = p < 0.01, ***=p<0.001, ****=p<0.0001, ns=p>=0.05.

## Results

### Cell cycle block by AURKA inhibitor but not PI3K inhibitor in SUM149 IBC cells

In a previous pharmacogenomics screening study, the cytotoxicity of pan-PI3K inhibitor Buparlisib in SUM149 IBC cells was significantly improved by silencing the *Aurka* gene (8). Here, we investigated whether pharmacological inhibition of AURKA using Alisertib would phenocopy the sensitizing effects to killing of IBC cells by Buparlisib. First, we tested the effects of Alisertib (ALS) and Buparlisib (BKM120) treatments alone or in combination (combo) on the PI3K pathway as indicated by phosphorylation of AKT (Fig. 1A). At a dose of 5 μM for each drug, treatments of SUM149 cells for 1 hour with Buparlisib alone, or in combination with Alisertib, led to greatly reduced AKT phosphorylation (pAKT) visualized by immunoblotting (Fig. 1A). As expected, Alisertib treatment alone did not alter AKT phosphorylation compared to DMSO vehicle control (Fig. 1A). Several experiments were analyzed by densitometry, and showed a significant reduction in relative levels of pAKT in Buparlisib and combo treatment groups (Fig. 1B). To investigate the effects of Alisertib treatments on AURKA activation, we attempted to measure phosphorylation of AURKA (pAURKA) in lysates from asynchronously growing SUM149 cells and failed to observe sufficient pAURKA signal (data not shown). However, using a double thymidine block to synchronize SUM149 cells in G1 phase, we released the cells for various time points to assess cell cycle status by DNA content analyzed by flow cytometry. We observed a high percentage of SUM149 cells in G2/M phase between 5 and 8 hours post release (Supplementary Figure 1). We used the double thymiding block and 7 hour release of SUM149 cells for testing effects of Alisertib on pAURKA, and observed a strong reduction compared to vehicle control (Fig. 1C/D). Buparlisib treatments did not impair pAURKA, but it was impaired in combination treatments with Alisertib (Fig. 1C/D). Thus, both inhibitors act on their target pathways in SUM149 cells and are compatible in combination treatments.

**Fig. 1.**
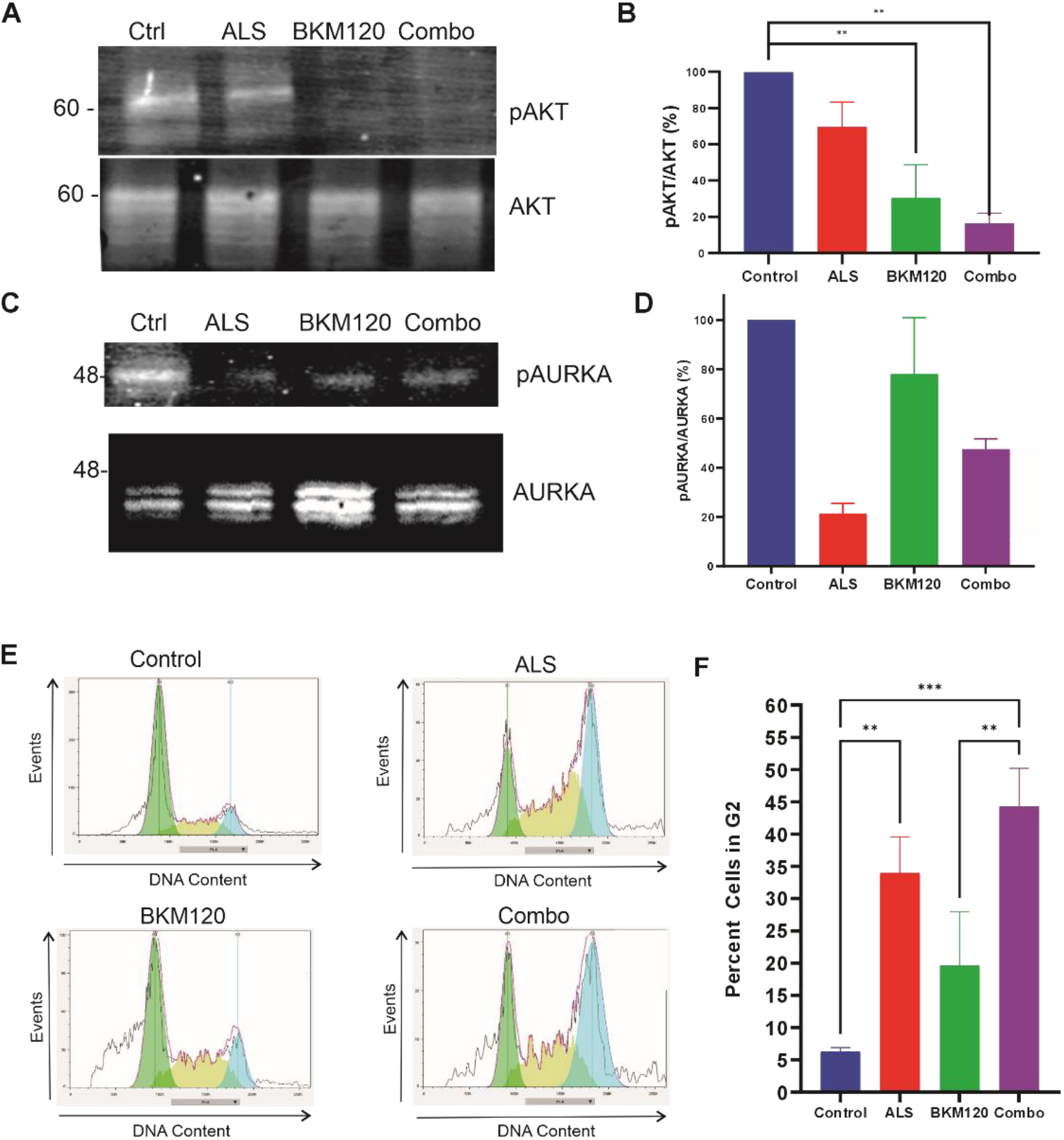
On-target effects of Alisertib and Buparlisib in SUM149 cells and cell cycle disruption by Alisertib. **a** SUM-149 cells were treated with DMSO vehicle control, Alisertib (ALS, 5 μM), Buparlisib (BMK120, 5 μM), and their combination (same doses) for 1 hour. Lysates were subjected to immunoblotting with phospho-AKT (pAKT) or pan-AKT (AKT) antibodies. **b** Bar graph shows the densitometry of the pAKT/AKT ratio using ImageJ with statistical analysis in GraphPad Prism (mean± SEM for N=3 experiments; ** p<0.01 based on ANOVA with multiple comparison testing). **c** Double thymidine blocked SUM-149 cells were released for 7 hours and treated with DMSO vehicle control, Buparlisib (5 μM), Alisertib (5 μM), and their combination (same doses). Lysates were subjected to immunoblotting with phospho-AURKA (pAURKA) or pan-AURKA (AURKA) antibodies. **d** Bar graph shows the densitometry of the pAKT/AKT ratio using ImageJ with statistical analysis in GraphPad Prism (mean± SEM for N=3 experiments). **e** SUM-149 cells were treated with Buparlisib or Alisertib alone (5 μM) or in combination (5 μM each) for 48 hours, and representative flow cytometry histograms are shown for Propidium Iodide-stained, permeabilized cells (DNA content). **f** Graph depicts the percentage of cells in G2 phase of the cell cycle from 3 experiments (mean ± SEM; ** p<0.01 or *** p<0.001 based on ANOVA with multiple comparison testing).

Since both PI3K and AURKA pathways can impact cell growth and cell cycle, we tested the effects of vehicle control, Alisertib or Buparlisib alone (5 μM), or in combination, on asynchronously growing SUM149 cells treated for 48 hours. Using propidium iodide (PI) staining of permeabilized cells from each treatment group, we analyzed the cell cycle profiles using flow cytometry. We observed a significant increase in percentage of SUM149 cells in G2/M with Alisertib treatment compared to the control and Buparlisib treatments (Fig. 1E). A similar block in G2/M was observed in combination treatments (Fig. 1E), and analysis of several experiments revealed that the G2/M block by Alisertib and combo were significant (Fig. 1F). These results showed that Alisertib treatments triggered a cell cycle arrest in SUM149 cells, distinct from the limited effects of Buparlisib alone on cell cycle.

### Buparlisib and Alisertib treatments caused synergistic killing of IBC cells

Using Alamar Blue as a metabolic indicator of SUM149 cell viability, we next performed dose response analyses with Buparlisib (BKM120) and Alisertib (ALS) treatments for 48 hours. Consistent with the relative resistance of SUM149 cells to Buparlisib, only a modest dose-dependent reduction in cell viability was observed with ̴ 70% viable cells at 10 μM dose (Fig. 2A). A similar dose-dependent reduction in SUM149 cell viability was observed with Alisertib (Fig. 2A). Next, we examined combinations of Buparlisib and Alisertib at each dose, and tested for synergistic effects by calculating the Bliss synergy score (15). We detected a robust synergy score of 36.117 for Buparlisib and Alisertib combination treatments on IBC SUM149 cells (Fig. 2B). At several doses of Alisertib and Buparlisib, the reduction in cell viability with the combination was significantly greater than either drug alone (Fig. 2C, Supplementary Fig. 2). Overall, these results demonstrated that pharmacological inhibition of AURKA can sensitize SUM149 IBC cells to killing by PI3K inhibitor Buparlisib, and this warranted further investigation in IBC models.

**Fig. 2.**
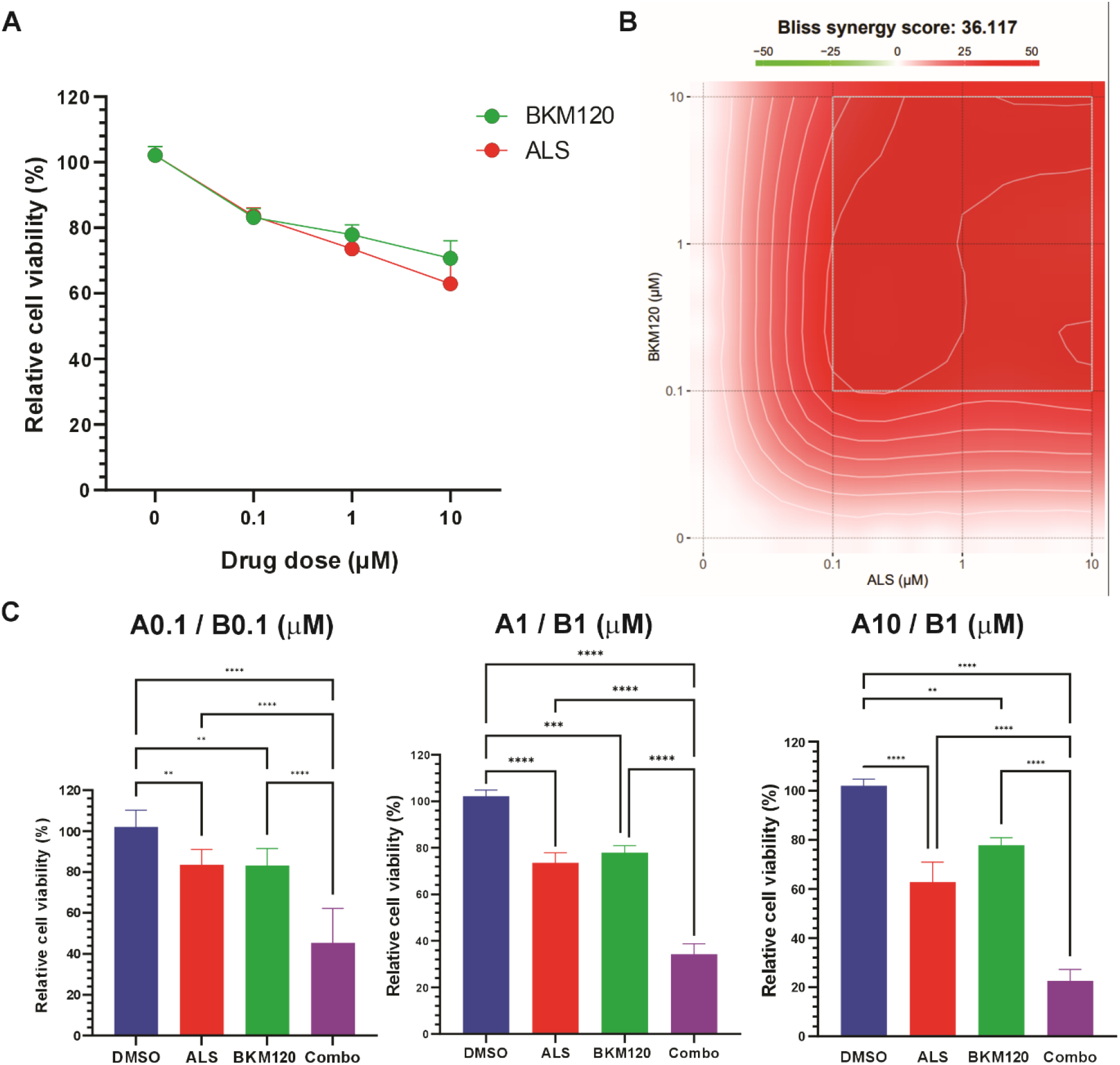
Synergistic killing of IBC cells by combination treatments with Buparlisib and Alisertib. **a** Relative cell viability of SUM149 cells was determined using Alamar blue assays at 48 hours post treatments with either Buparlisib (10, 1, or 0.1 μM) or Alisertib (10, 1, and 0.1 μM), compared to vehicle control (shown as 0 on the x-axis; mean ± SEM). **b** Plot represents the Bliss synergy score for combinations of Alisertib (ALS) and Buparlisib (BKM120) at multiple doses (10, 1, or 0.1 μM) on cell viability determined using Alamar blue at 48 hours. **c** Bar graphs depict relative cell viability at 48 hours for particular doses of Alisertib and Buparlisib at 0.1 μM (A0.1/B0.1), 1 μM (A1/B1), and at a 10:1 ratio of Alisertib:Buparlisib (A10/B1) as measured by Alamar blue. The results were obtained from three independent experiments, each performed in triplicate. The data were analyzed using GraphPad Prism and SynergyFinder software (detailed in Methods; mean ± SEM; ** p<0.01 or *** p<0.001 or **** p<0.0001 based on ANOVA with multiple comparison testing).

### Buparlisib and Alisertib treatments caused synergistic reduction of anchorage-independent growth and cell migration of IBC cells

Anchorage-independent growth of cancer cells has been used to test effects of genes or drugs on the transformed phenotype and resistance to anoikis. Here, SUM149 cells were seeded in soft agar and treated with vehicle control, Alisertib (5 μM) or Buparlisib (2.5 μM) alone, or in combination for 21 days (media was supplemented with drugs every 2 days). At endpoint, the colonies were stained with Crystal Violet and imaging revealed overt differences in some treatment groups compared to control (Fig. 3A). Alisertib treatments reduced colonies compared to controls, whereas Buparlisib treatments had no effect (Fig. 3B). Importantly, the reduction in colonies was greatest in the combination treatments, showing strong synergistic effects of Alisertib and Buparlisib in this relatively long term assay of IBC cell growth.

**Fig. 3.**
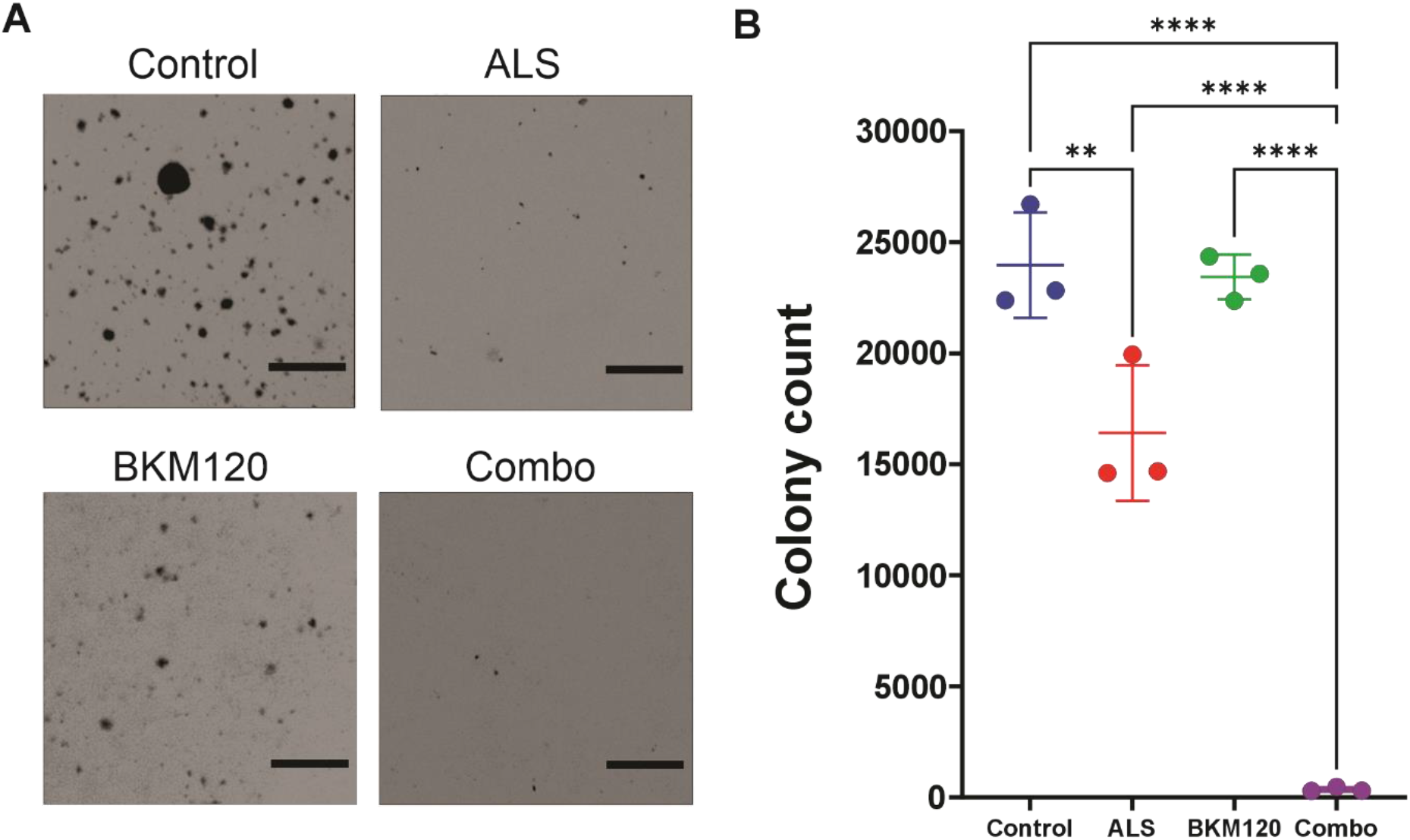
Synergistic effects of Buparlisib and Alisertib limiting anchorage-independent growth of IBC cells. **a** SUM149 cells were subjected to soft agar colony assays in presence of vehicle control, Alisertib (ALS, 5μM), Buparlisib (2.5 μM), or the combination at preceding doses (combo) for 21 days with drugs added fresh every 2 days. Representative micrographs of crystal violet-stained wells are shown (scale bar indicates 100 μm). **b** Graph depicts the scoring of average colony numbers per treatment group from 3 independent experiments (mean ± SEM; ** p<0.01 or **** p<0.0001 based on ANOVA with multiple comparison testing).

Next, we tested the effects of Alisertib and Buparlisib treatments on SUM 149 cell motility using wound healing assays. The cells were seeded at confluence in a 96-well plate, and a scratch wound was created prior to applying drug treatments or vehicle control. Phase-contrast images were captured every 2 hours for 24 hours. Representative images of the wound area at time 0 or 24 hours show that SUM149 cells completely close the wound area with control, but less so with Alisertib, Buparlisib or combination treatments (Fig. 4A). Quantification of the percentage wound confluence was calculated for each treatment group, and showed reduced motility with Alisertib or Buparlisib alone compared to control (Fig. 4B). Importantly, the wound healing migration was reduced most significantly by the combination treatments (Fig. 4B). In summary, these results demonstrated that the combination of Alisertib and Buparlisib significantly impaired cell migration in SUM149 cells *in vitro*, and this may help restrain the metastatic properties of IBC *in vivo*.

**Fig. 4.**
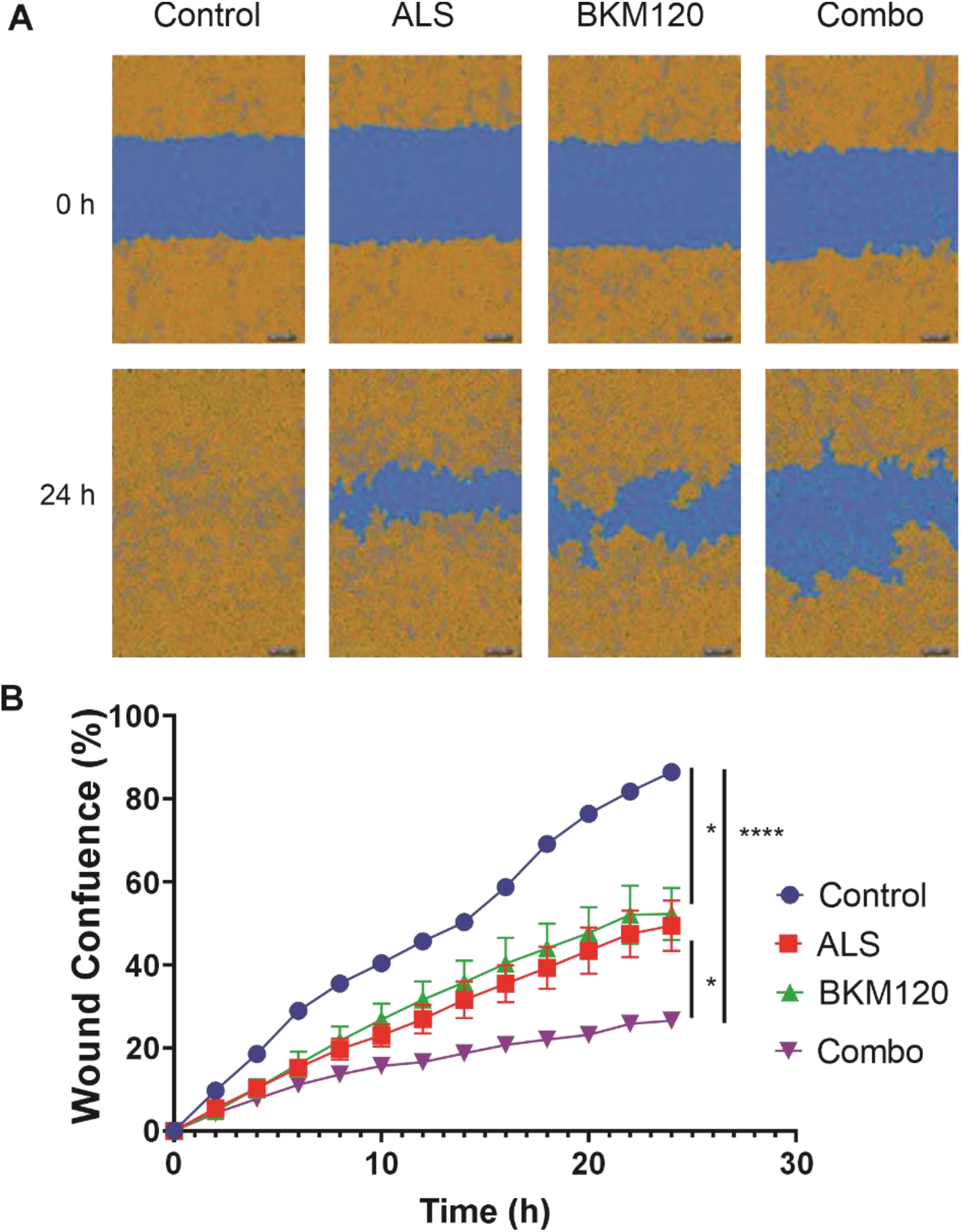
Alisertib and Buparlisib treatments reduce IBC cell motility. **a** SUM149 cell motility was analyzed using scratch wound assays using an 96 well plate wound maker and Incucyte Zoom system (Sartorius). Treatment groups included vehicle control, Alisertib (ALS, 0.5 μM), Buparlisib (BKM120, 0.5 μM), or the combination at preceding doses over 24 hours. Representative images are shown for time 0 and 24 hours of the wound area mask (blue) and cell mask (yellow) for each treatment group. **b** The graph illustrates the increase in percent wound confluence rates for each treatment group (mean ± SEM; * p<0.05 or **** p<0.0001 based on ANOVA with multiple comparison testing). The results are representative of 3 independent experiments.

### Reduced tumor growth and metastasis in IBC tumor-bearing mice treated with Alisertib and Buparlisib

To extend studies of Alisertib and Buparlisib treatments to tumor growth and metastasis, we performed mammary orthotopic tumor xenograft assays with SUM149 cells implanted within immunocompromised female mice (Rag2^-/-^:IL2Rγc^-/-^). When palpable tumors were detected, mice were randomized into 4 different groups with 6 mice per group on day 12. The control group received intraperitoneal (i.p.) injections with the vehicle used to solubilize both drugs for in vivo studies. Both Alisertib (ALS) and Buparlisib (BKM120) were dosed at 25 mg/kg (i.p.; once daily) as either monotherapies or combination therapy with a 5 days on and 2 days off treatment schedule (5 + 2; Fig. 5A). After 35 days, the animals were culled, and tumors and tissues were harvested for further processing. Significant reductions in tumor weights were observed in Alisertib and combination treatment groups (Fig. 5B). Upon histological analysis of tumor tissue sections, we observed necrotic areas within the tumors from the Alisertib and Buparlisib or combo treatment groups (Fig. 5C; annotated by areas labeled N). Quantification of the percent necrosis area showed an increasing trend in the combination group, and thus showed the most promising control of IBC tumor growth in this IBC xenograft model.

**Fig. 5.**
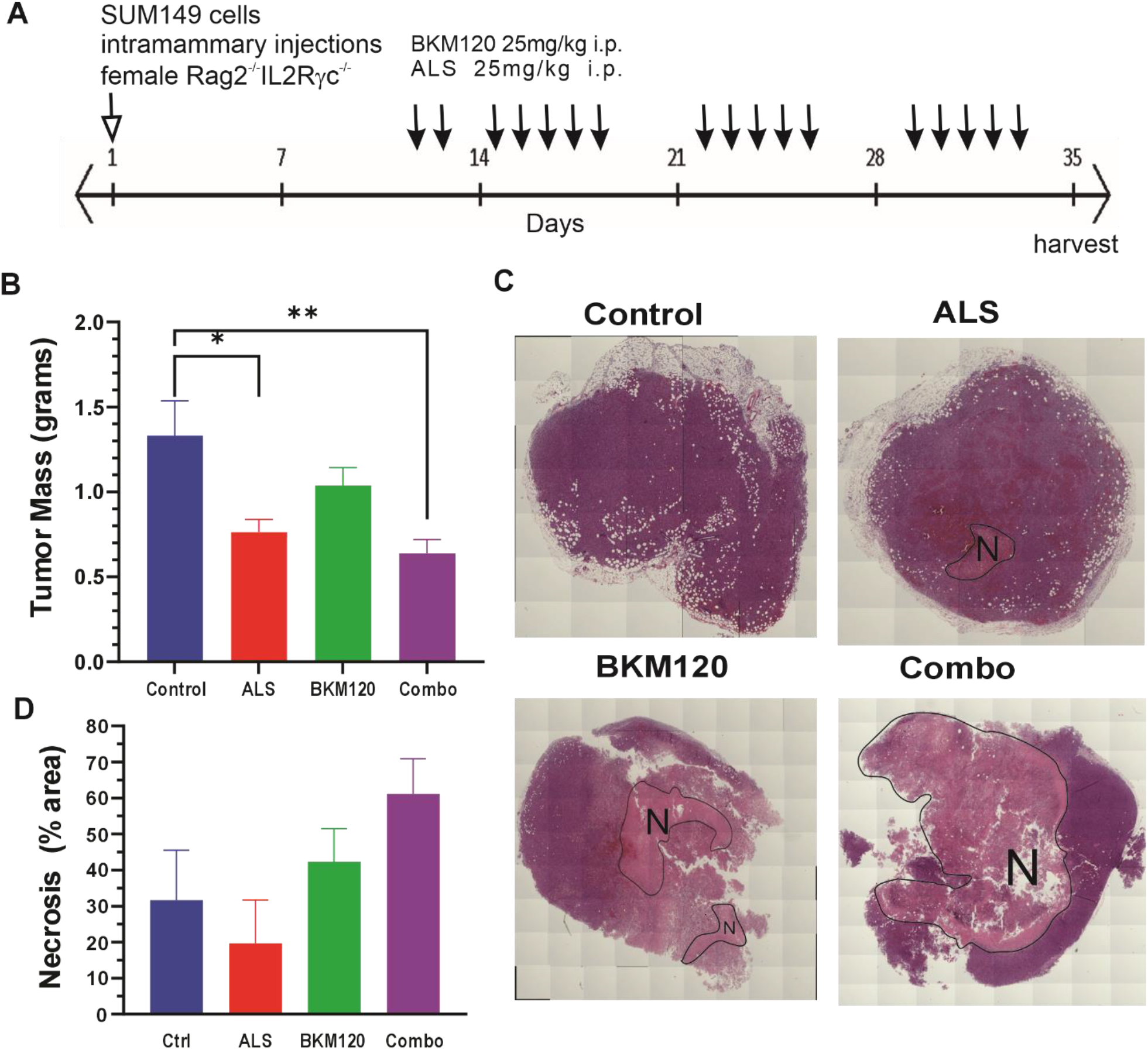
Buparlisib and Alisertib inhibit IBC tumor growth *in vivo*. **a** A timeline of the mammary orthotopic SUM149 tumor xenograft assays using female Rag2^-/-^:IL2R ^-/-^ mice. Mice were split into four groups (6 mice per group) on day 12 and treated with vehicle control, Buparlisib (BKM120, 25 mg/kg), Alisertib (ALS, 25 mg/kg), and combination treatment by intraperitoneal injections on a 5 on and 2 off schedule. **b** Graph depicts tumor mass measured at endpoint (day 35, mean ± SEM, * p<0.05 or ** p<0.01 based on ANOVA with multiple comparison testing). **c** Representative images showing H&E-stained tumor tissue sections (black freehand shape depicts areas of necrosis). **d** The graph depicts the percentage area of necrosis within tumor cross-sections by treatment group (mean ± SEM).

We next analyzed the lung tissues from the above mammary orthotopic IBC xenograft study to analyze treatment effects on spontaneous metastases to the lungs of these mice. We stitched together phase contrast images of hematoxylin/eosin-stained lung tissue sections spanning an entire lobe of the lung from all the animals. Representative images from each treatment group showed areas with micrometastases (Fig. 6A, see high magnification inserts). Scoring of these metastases was performed by two independent investigators, and the average numbers of metastases were determined for each animal (Fig. 6B). Treatments with Alisertib or Buparlisib alone reduced the frequency of lung metastases, with the fewest detected within the combination treatment group (Fig. 6B). Together, these results support the potential benefits of treating IBC with Alisertib and Buparlisib combination therapy to restrain both tumor growth and metastases.

**Fig. 6.**
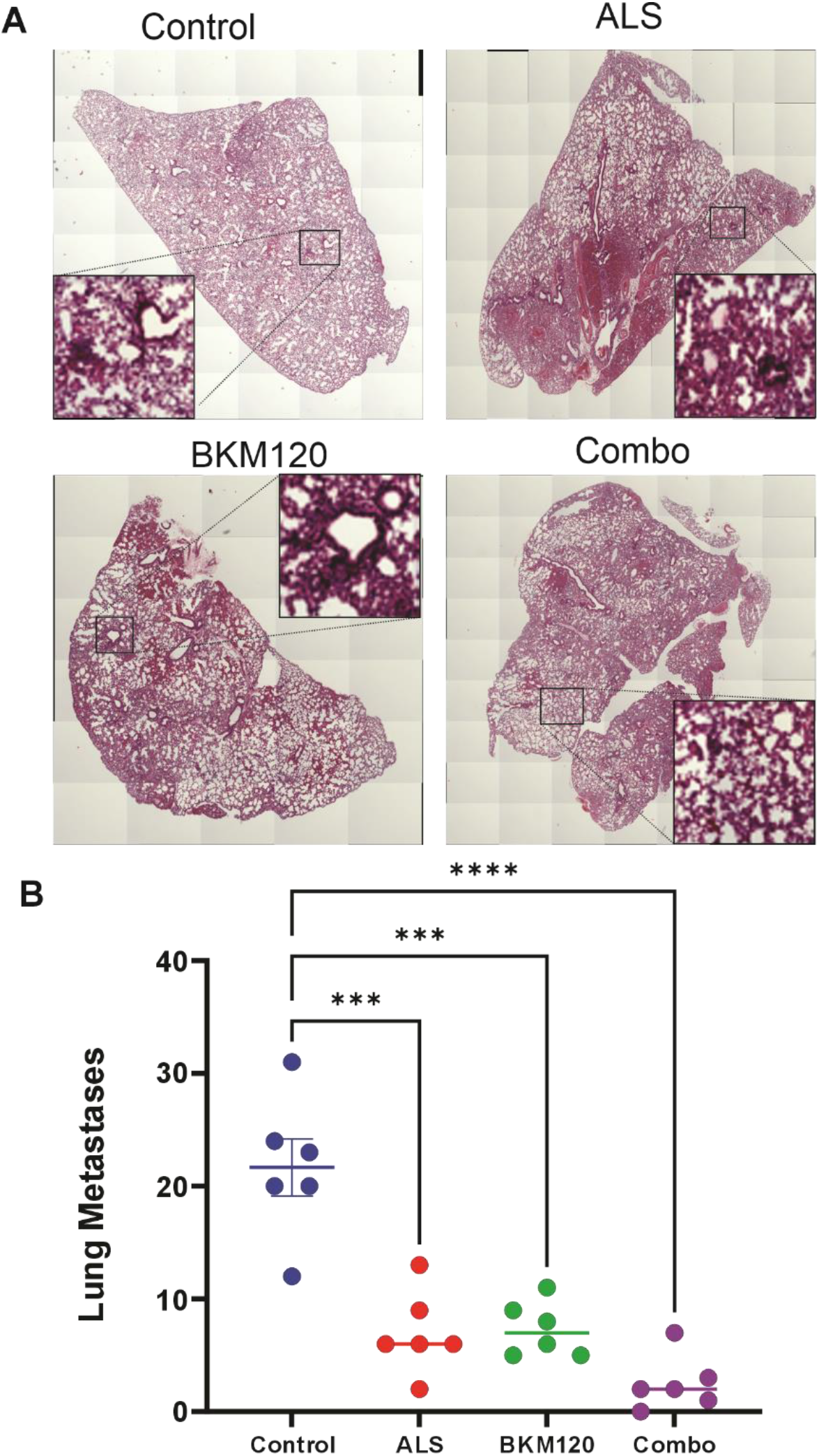
Reduced IBC lung metastasis in Buparlisib and Alisertib treated mice. **a** Representative images showing H&E-stained lung tissue sections from each treatment group. The areas within the black squares are shown at higher magnification insets for each treatments group. **b** Graph depicts the average scoring of lung metastases for each animal in the mammary orthotopic IBC xenograft model for each treatment group (mean ± SEM, *** p<0.001 or **** p<0.0001 based on ANOVA with multiple comparison testing).

## Discussion

In this study, we provide rationale and evidence that inhibiting AURKA can sensitize IBC cells and tumors to killing by the pan-PI3K inhibitor Buparlisib. Our findings address the limited targeted therapy options for IBC (16), and the need to identify additional pathways that provide resistance to PI3K inhibitors (8). The PI3K pathway is hyperactivated in almost all breast cancer types, including IBC (17),(18), including gain-of-function hot spot mutations in PIK3CA and deletions of the PTEN tumor suppressor gene (19). Dysregulation in the PI3K pathway hinders the action of several anticancer drugs, which led to the development of several PI3K inhibitors (8),(17),(18). This study focused on the AURKA pathway as a potential co-target with the PI3K pathway since Aurka gene silencing was shown to enhance killing of IBC cells by Buparlisib (8). The AURKA pathway plays a crucial role in cell cycle progression, particularly during mitosis (20). Cross-talk between the AURKA pathway as also been observed with other oncogenic pathways (21). We chose AURKA inhibitor Alisertib to test in combination with Buparlisib since it has shows some efficacy in breast cancer cell lines and modest toxicities (21). Alisertib treatments disrupt key processes in mitosis, including chromosome alignment and spindle bipolarity (22). This is consistent with our findings that IBC cells accumulate in G2 phase upon treatments with Alisertib, and this may explain synergy with Buparlisib in killing these checkpoint restricted cells.

While Buparlisib has shown efficacy in killing IBC cell lines, clinical trials has shown limited efficacy as monotherapy and raised concerns of toxicity profiles (20),(23),(24),(25). We suggest exploring alternative, class-specific PI3K inhibitors such as Alpelisib in future studies. In fact, Alpelisib has demonstrated efficacy in targeting PIK3CA-mutated cancer models and has received FDA approval for patients with advanced ER-positive breast cancer (26),(27). It will be interesting to test for synergy between Alpelisib and Alisertib in both IBC and non-IBC cell and tumor models in future.

Metastasis is a significant concern in IBC, particularly with lung involvement (28),(29). In this study, we show that the combination of Alisertib and Buparlisib inhibited cell migration of IBC cells in vitro, and reduced lung metastases in IBC tumor xenograft assays. We observed more areas of necrosis within the tumors from mice treated with the combination of Buparlisib and Alisertib. Future studies will be needed to understand if treatments impact local invasion in the mammary tissue, killing of the lung-resident metastatic cells, or both. Furthermore, recent studies highlight the involvement of extrinsic factors and genes controlling epithelial-mesenchymal plasticity, migration, and invasion in IBC (19),(30),(31),(32). Since the siRNA screening study of kinase genes that are synthetically lethal with PI3K inhibitor treatment in IBC cells (8), it would be interesting to extend these studies to genome-wide screens using CRISPR/Cas9 approaches.

## Conclusions

Our study provides evidence supporting efficacy of a combination therapy comprising Buparlisib and Alisertib to treat IBC. By concurrently targeting two pivotal pathways, namely PI3K and AURKA, our approach effectively disrupts the oncogenic addiction of IBC cells to these pathways. The strong synergistic effects of Buparlisib and Alisertib may overcome resistance mechanisms to either inhibitor as a monotherapy. Further testing in additional models will likely be needed to advance these findings to future human clinical trials in IBC.

## Abbreviation List

IBC: Inflammatory breast cancer
ER/PR/HER2: oestrogen/progesterone/ human epidermal growth factor 2 receptors
PI3K: phosphoinositide-3-kinase
AURKA: Aurora kinase A
BKM120: Buparlisib
ALS: Alisertib
DMSO: Dimethyl sulfoxide.

## Availability of Data and Material

The datasets used and/or analysed during the current study are available from the corresponding author upon reasonable request.

## Funding

This research was funded by Queen’s University Research Opportunity Fund grant and by a grant from Canadian Institutes of Health Research to AWC. Salary support for NAA and JK was provided by Queen’s Graduate awards and Queen’s Health Sciences Dean’s award.

## Acknowledgements

The authors thank other Craig lab members for providing helpful advice and support during the course of this research.

## Declarations

### Ethical Approval

Research with human cancer cell lines and animals were approved by the Queen’s University Biohazard and Animal Care committees, respectively.

### Competing interests

The authors declare no financial or personal competing interests related to this research.

### Authors’ contributions

NAA, JK, and SY performed the experiments and analyzed the results. NAA wrote the initial draft of the manuscript. AWBC conceived of the study, helped interpret the results and edited the manuscript.

## Notes

### Competing Interest Statement

The authors have declared no competing interest.

## References

1. Devi GR, Hough H, Barrett N, Cristofanilli M, Overmoyer B, Spector N, et al. Perspectives on Inflammatory Breast Cancer (IBC) Research, Clinical Management and Community Engagement from the Duke IBC Consortium. J Cancer. 2019;10(15):3344–51.

2. Costa R, Santa-Maria CA, Rossi G, Carneiro BA, Chae YK, Gradishar WJ, et al. Developmental therapeutics for inflammatory breast cancer: Biology and translational directions. Oncotarget. 2017;8(7):12417–32.

3. Mamouch F, Berrada N, Aoullay Z, El Khanoussi B, Errihani H. Inflammatory Breast Cancer: A Literature Review. World J Oncol. 2018;9(5-6):129–35.

4. Liang X, Vacher S, Boulai A, Bernard V, Baulande S, Bohec M, et al. Targeted next-generation sequencing identifies clinically relevant somatic mutations in a large cohort of inflammatory breast cancer. Breast Cancer Res. 2018;20(1):88.

5. Yang J, Nie J, Ma X, Wei Y, Peng Y, Wei X. Targeting PI3K in cancer: mechanisms and advances in clinical trials. Mol Cancer. 2019;18(1):26.

6. Vagia E, Mahalingam D, Cristofanilli M. The Landscape of Targeted Therapies in TNBC. Cancers (Basel). 2020;12(4).

7. Wright SCE, Vasilevski N, Serra V, Rodon J, Eichhorn PJA. Mechanisms of Resistance to PI3K Inhibitors in Cancer: Adaptive Responses, Drug Tolerance and Cellular Plasticity. Cancers (Basel). 2021;13(7).

8. Van Swearingen AED, Sambade MJ, Siegel MB, Sud S, McNeill RS, Bevill SM, et al. Combined kinase inhibitors of MEK1/2 and either PI3K or PDGFR are efficacious in intracranial triple-negative breast cancer. Neuro Oncol. 2017;19(11):1481–93.

9. Mou PK, Yang EJ, Shi C, Ren G, Tao S, Shim JS. Aurora kinase A, a synthetic lethal target for precision cancer medicine. Exp Mol Med. 2021;53(5):835–47.

10. Courtheoux T, Diallo A, Damodaran AP, Reboutier D, Watrin E, Prigent C. Aurora A kinase activity is required to maintain an active spindle assembly checkpoint during prometaphase. J Cell Sci. 2018;131(7).

11. Damodaran AP, Vaufrey L, Gavard O, Prigent C. Aurora A Kinase Is a Priority Pharmaceutical Target for the Treatment of Cancers. Trends Pharmacol Sci. 2017;38(8):687–700.

12. Manfredi MG, Ecsedy JA, Chakravarty A, Silverman L, Zhang M, Hoar KM, et al. Characterization of Alisertib (MLN8237), an investigational small-molecule inhibitor of aurora A kinase using novel in vivo pharmacodynamic assays. Clin Cancer Res. 2011;17(24):7614–24.

13. Cervantes A, Elez E, Roda D, Ecsedy J, Macarulla T, Venkatakrishnan K, et al. Phase I pharmacokinetic/pharmacodynamic study of MLN8237, an investigational, oral, selective aurora a kinase inhibitor, in patients with advanced solid tumors. Clin Cancer Res. 2012;18(17):4764–74.

14. Ianevski A, Giri AK, Aittokallio T. SynergyFinder 2.0: visual analytics of multi-drug combination synergies. Nucleic Acids Res. 2020;48(W1):W488–W93.

15. Buchheit CL, Angarola BL, Steiner A, Weigel KJ, Schafer ZT. Anoikis evasion in inflammatory breast cancer cells is mediated by Bim-EL sequestration. Cell Death Differ. 2015;22(8):1275–86.

16. Chainitikun S, Espinosa Fernandez JR, Long JP, Iwase T, Kida K, Wang X, et al. Pathological complete response of adding targeted therapy to neoadjuvant chemotherapy for inflammatory breast cancer: A systematic review. PLoS One. 2021;16(4):e0250057.

17. Criscitiello C, Viale G, Curigliano G, Goldhirsch A. Profile of buparlisib and its potential in the treatment of breast cancer: evidence to date. Breast Cancer (Dove Med Press). 2018;10:23–9.

18. Du R, Huang C, Liu K, Li X, Dong Z. Targeting AURKA in Cancer: molecular mechanisms and opportunities for Cancer therapy. Mol Cancer. 2021;20(1):15.

19. Carbognin L, Miglietta F, Paris I, Dieci MV. Prognostic and Predictive Implications of PTEN in Breast Cancer: Unfulfilled Promises but Intriguing Perspectives. Cancers (Basel). 2019;11(9).

20. Nikonova AS, Astsaturov I, Serebriiskii IG, Dunbrack RL, Golemis EA. Aurora A kinase (AURKA) in normal and pathological cell division. Cell Mol Life Sci. 2013;70(4):661–87.

21. Jalalirad M, Haddad TC, Salisbury JL, Radisky D, Zhang M, Schroeder M, et al. Aurora-A kinase oncogenic signaling mediates TGF-β-induced triple-negative breast cancer plasticity and chemoresistance. Oncogene. 2021;40(14):2509–23.

22. Niu H, Manfredi M, Ecsedy JA. Scientific Rationale Supporting the Clinical Development Strategy for the Investigational Aurora A Kinase Inhibitor Alisertib in Cancer. Front Oncol. 2015;5:189.

23. Patsouris A, Augereau P, Frenel JS, Robert M, Gourmelon C, Bourbouloux E, et al. Benefits versus risk profile of buparlisib for the treatment of breast cancer. Expert Opin Drug Saf. 2019;18(7):553–62.

24. Garrido-Castro AC, Saura C, Barroso-Sousa R, Guo H, Ciruelos E, Bermejo B, et al. Phase 2 study of buparlisib (BKM120), a pan-class I PI3K inhibitor, in patients with metastatic triple-negative breast cancer. Breast Cancer Res. 2020;22(1):120.

25. Xing J, Yang J, Gu Y, Yi J. Research update on the anticancer effects of buparlisib. Oncol Lett. 2021;21(4):266.

26. Narayan P, Prowell TM, Gao JJ, Fernandes LL, Li E, Jiang X, et al. FDA Approval Summary: Alpelisib Plus Fulvestrant for Patients with HR-positive, HER2-negative, PIK3CA-mutated, Advanced or Metastatic Breast Cancer. Clin Cancer Res. 2021;27(7):1842–9.

27. André F, Ciruelos EM, Juric D, Loibl S, Campone M, Mayer IA, et al. Alpelisib plus fulvestrant for PIK3CA-mutated, hormone receptor-positive, human epidermal growth factor receptor-2-negative advanced breast cancer: final overall survival results from SOLAR-1. Ann Oncol. 2021;32(2):208–17.

28. van Uden DJP, van Maaren MC, Strobbe LJA, Bult P, van der Hoeven JJ, Siesling S, et al. Metastatic behavior and overall survival according to breast cancer subtypes in stage IV inflammatory breast cancer. Breast Cancer Res. 2019;21(1):113.

29. Dano D, Lardy-Cleaud A, Monneur A, Quenel-Tueux N, Levy C, Mouret-Reynier MA, et al. Metastatic inflammatory breast cancer: survival outcomes and prognostic factors in the national, multicentric, and real-life French cohort (ESME). ESMO Open. 2021;6(4):100220.

30. Kvokačková B, Remšík J, Jolly MK, Souček K. Phenotypic Heterogeneity of Triple-Negative Breast Cancer Mediated by Epithelial-Mesenchymal Plasticity. Cancers (Basel). 2021;13(9).

31. Li JJ, Sun ZJ, Yuan YM, Yin FF, Bian YG, Long LY, et al. EphB3 Stimulates Cell Migration and Metastasis in a Kinase-dependent Manner through Vav2-Rho GTPase Axis in Papillary Thyroid Cancer. J Biol Chem. 2017;292(3):1112–21.

32. Yang Z, He J, Gao P, Niu Y, Zhang J, Wang L, et al. miR-769-5p suppressed cell proliferation, migration and invasion by targeting TGFBR1 in non-small cell lung carcinoma. Oncotarget. 2017;8(69):113558–70.

