## Supplemental materials and figures for "Synergistic effects of inhibitors targeting PI3K and Aurora Kinase A in preclinical inflammatory breast cancer models"

**Supplementary materials**

The original immunoblots from Fig. 1A and Fig. 1C are provided below.

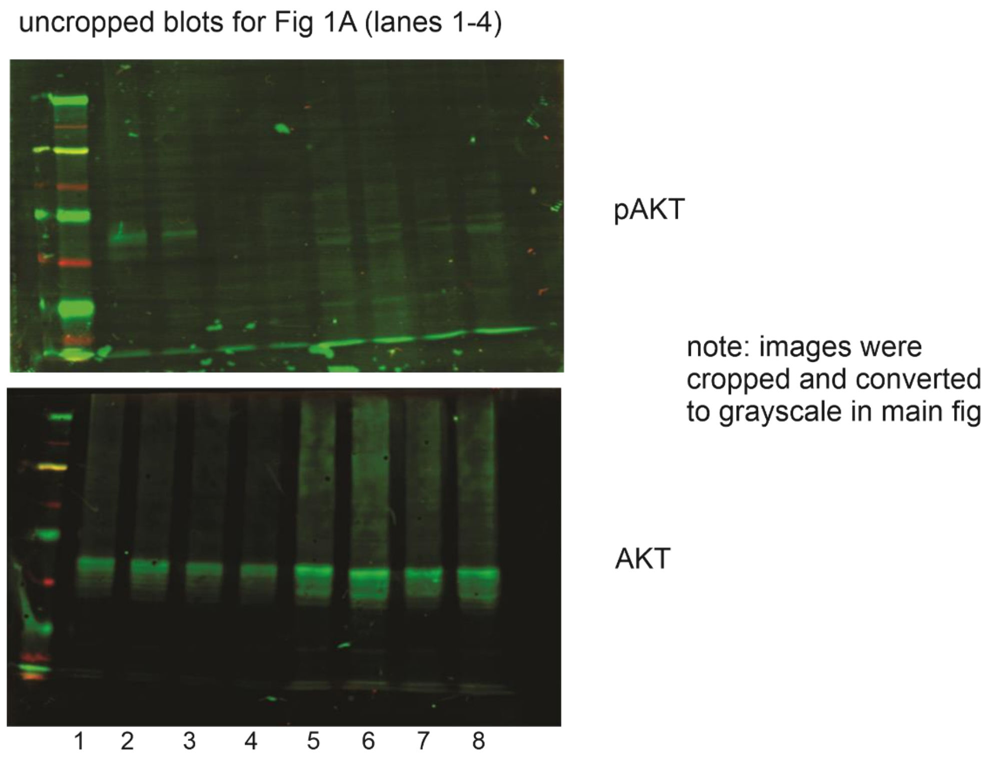


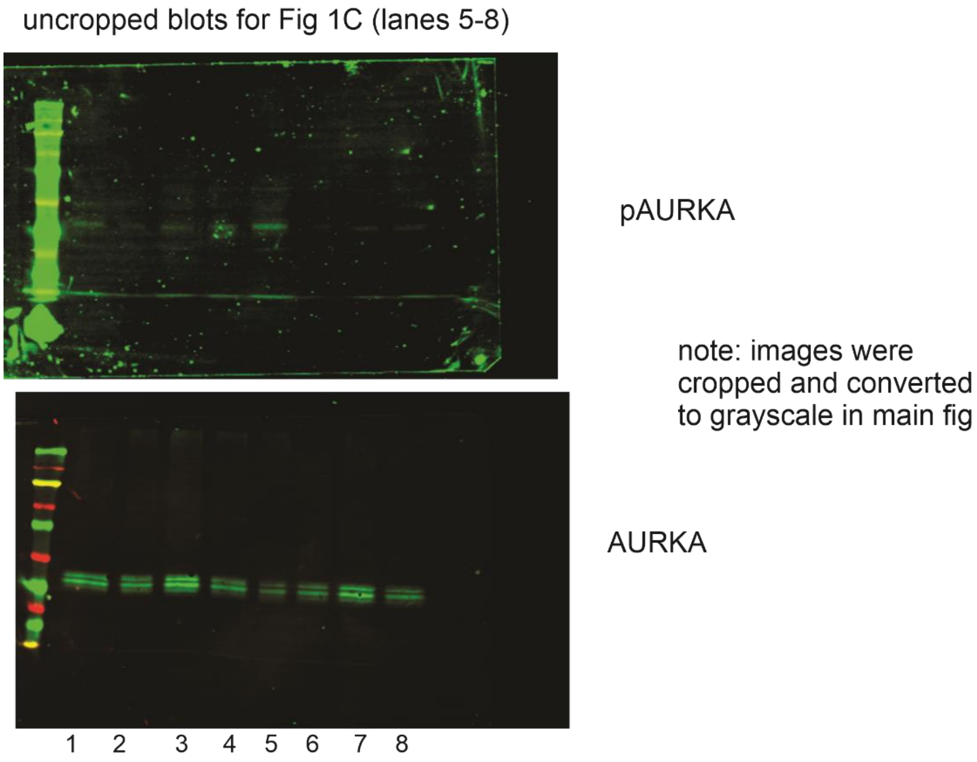

### Supplementary Figures

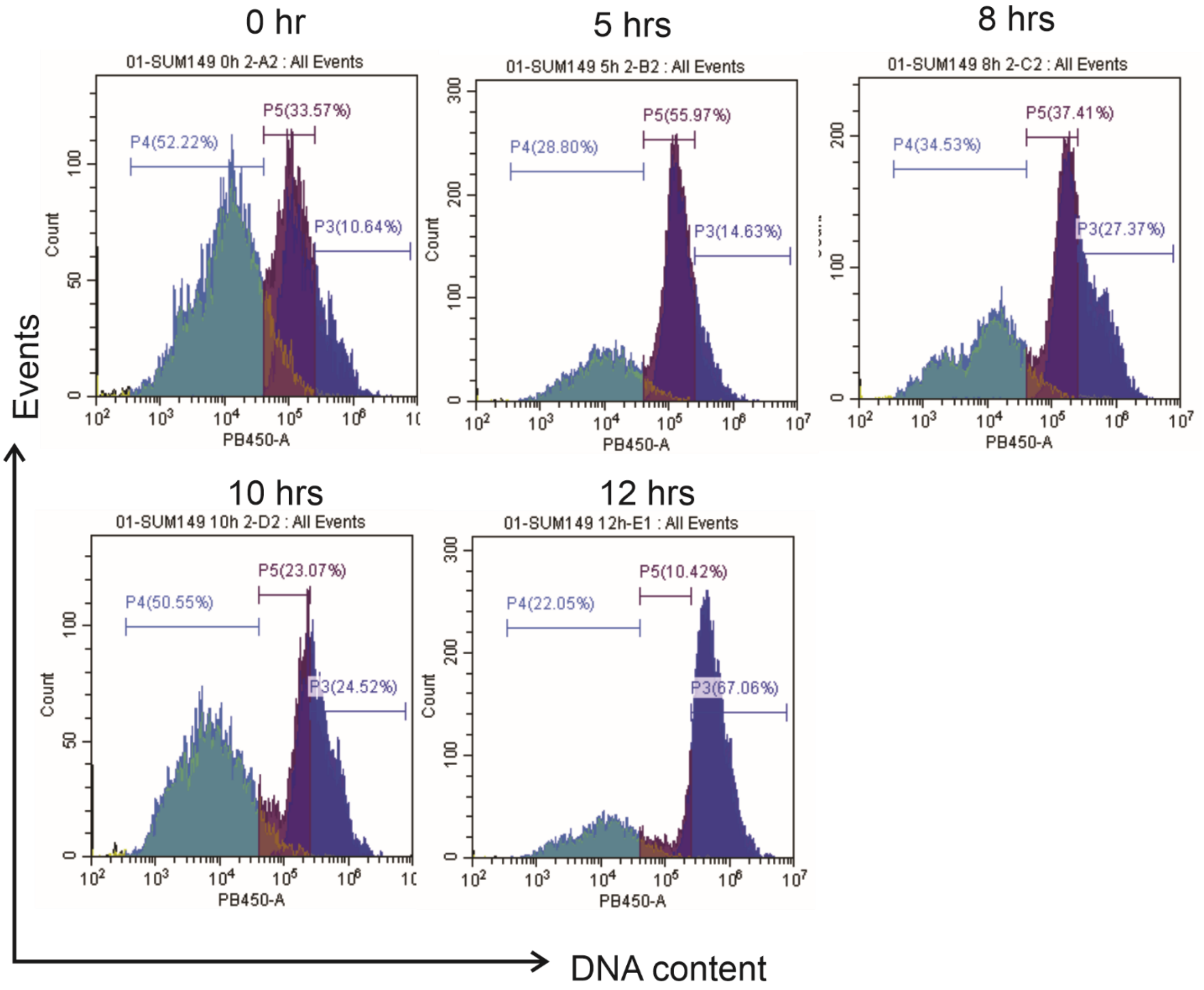


**Supplementary Fig. 1** Double thymidine block and release of synchronized SUM-149 cells to assess cell cycle. Cells at the indicated times post release in fresh media were collected, permeabilized and stained with PI for analysis of DNA content by flow cytometry. Representative histograms are shown.

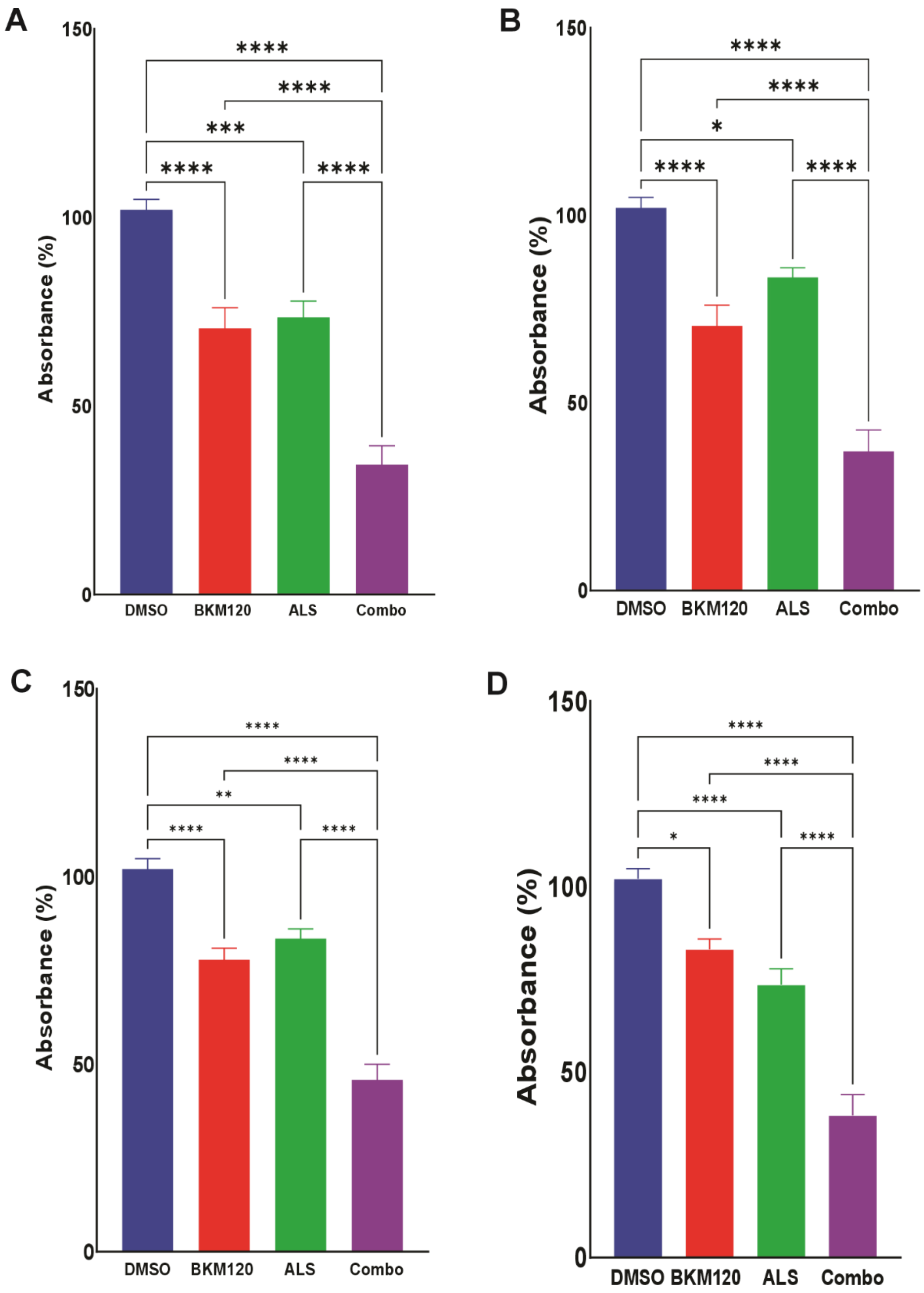


**Supplementary Fig. 2** SUM149 cell viability profiles comparing various doses of Buparlisib and Alisertib treatments. **a** Endpoint (48 hour) cell viability data plotted from the Alamar blue assays with SUM149 cells treated with BKM120 and/or ALS at 10 and 1 μM, respectively. **b** Endpoint (48 hour) cell viability data plotted from Alamar blue assays with SUM149 cells treated with BKM120 and/or ALS at 10 and 0.1 μM, respectively. **c** Endpoint (48 hour) cell viability data plotted from the Alamar blue assay with SUM149 cells treated with BKM120 and/or ALS at 1 and 0.1 μM, respectively. **d** Endpoint (48 hour) cell viability data plotted from Alamar blue assay with SUM149 cells treated with BKM120 and/or ALS at 0.1 and 1 μM, respectively. ANOVA with multiple comparison testing was performed to identify statistically significant differences between treatments for repeated experiments (*$p<0.05$, **$p<0.01$, ***$p<0.001$, ****$p<0.0001$).
